# Electrical Potentials of Protoscoleces of the Cestode *Echinococcus granulosus* from Bovine Origin

**DOI:** 10.1101/2021.06.16.448647

**Authors:** Mónica P. A. Carabajal, Marcos A. Durán, Santiago Olivera, María José Fernández Salom, Horacio F. Cantiello

## Abstract

Larval stages of the tenia *Echinococcus granulosus* are the infective forms of cystic echinococcosis or hydatidosis, a worldwide zoonosis. The protoscolex that develops into the adult form in the definitive host is enveloped by a complex cellular syncytial tegument, where all metabolic interchange takes place. Little information is available as to the electrical activity of the parasite in this developmental stage. To gain insight into the electrical activity of the parasite at the larval stage, here we conducted microelectrode impalements of bovine lung protoscoleces (PSCs) of *Echinococcus granulosus* in normal saline solution. We observed two distinct intra-parasitic potentials, a transient peak potential and a stable second potential, most likely representing tegumental and intra-parasitic extracellular space electrical potential differences, respectively. These values changed upon the developmental status of the parasite, its anatomical regions, or time course after harvesting. Changes in electrical potential differences of the parasite provide an accessible and useful parameter for the study of transport mechanisms and potential targets for the development of novel antiparasitic therapeutics.

**Author summary:** Hydatid disease is a parasite-caused zoonosis that causes high morbidity and mortality and has a great impact on public health. The disease has no known cure, and the main lines of treatment include surgery and medical treatments which are not satisfactory, so new drug compounds are urgently needed. Genome sequencing of the parasite has identified different genes encoding ion channels in *Echinococcus granulosus*, making ion channel inhibitors a promising target for treating hydatidosis. However, no easy technical approaches are available to test the electrical contribution of ion channels to parasite physiology. In the present study, we used the microelectrode impalement technique to determine the electrical properties of the larval stages of the parasite harvested from infected cow lungs. We observed transient electrical potentials not previously reported for the parasite, and changes in these parameters associated with its developmental stage and aging. Our findings indicate that microelectrode impalement of protoscoleces of *Echinococcus granulosus* may be a strategy of choice to explore and test possible drugs suggested for their therapeutic potential against hydatid disease. Further evaluation of parasites coming from other animals and humans may help address important issues in the treatment and prevention of the hydatid disease.

## Introduction

Cystic echinococcosis or hydatidosis encompasses a group of important zoonotic diseases caused by the metacestode (larval stage) of Taenia tapeworms belonging to the genus *Echinococcus*. The disease is mainly transmitted in livestock-raising areas, and the two most important zoonotic species of this genus are *E. granulosus* and *E. multilocularis*, causing cystic -or unilocular-echinococcosis (CE) and alveolar -or multivesicular-echinococcosis (AE), respectively [1, 2]. *E. granulosus* is the most widespread, with endemic foci in various continents, including South America, the entire Mediterranean littoral, central Asia, China, Australia, and Africa. In South America, CE is endemic in Argentina, Uruguay, Brazil, Chile, and some regions of Perú and Bolivia [3]. In Argentina, this zoonosis spreads through genetically distinct populations of the parasite [4].

*E. granulosus* undergoes a developmental cycle where sexual development of the adult stage occurs in the small intestine of dogs. Excreted eggs (larval stage) undergo embryonic development into oncospheres that scatter to the environment by fecal deposition. The cycle continues after ingestion of oncospheres by intermediate hosts, including humans. The embryos that hatch from the eggs penetrate the intestinal mucosa and distribute to the liver or other organs, undergoing metamorphosis into the next larval stage, the metacestode [3]. Closing the life-cycle, the metacestode is eventually ingested by a definitive host (mostly dog) to develop into a segmented and sexually mature adult stage again [5].

Metacestodes constitute fluid-filled cysts with an inner thin germinal layer where large numbers of protoscoleces (PSCs) are formed by asexual multiplication. The germinal layer invaginates to form vesicles and brood capsules [6]. PSCs remain invaginated within the mucopolysaccharide-coated basal region of the protoscoleces tegument (invaginated PSC) to protect the scolex until evagination in the definitive host (evaginated PSC). The precise stimulation for parasite evagination remains unknown, but environmental changes such as variations in temperature and osmotic pressure may be among the determining factors [6, 7].

Like other cestodes, *E. granulosus* lacks a digestive system, such that the parasite absorbs water, salts, and nutrients through the external tegument [8]. Thus, the study of the functional properties of the syncytial tegument is necessary for understanding host-parasite interactions. However, little information is available about the physiological differences between the invagination/evagination processes in protoscoleces (PSCs) of *E. granulosus*. Electrophysiological techniques are useful in the assessment of the electrical properties of PSCs and can help in the understanding of electrolyte transport by the tegumental epithelium. Microelectrode recordings have previously been used to characterize tegumental potential differences (PD) of *Schistosoma mansoni* [9] and the tegumental potential differences (PD) of ovine strain *E. granulosus* [8], where significant changes were observed upon immunological and chemical manipulations [10, 11].

Here we used microelectrode recordings from PSCs of *E. granulosus* from lungs of bovine origin to obtain parasitic potential differences (PD). Two distinct parasitic PD, transient and stable trans-tegumental PD, were recorded from both invaginated and evaginated PSCs. Data provide, to our knowledge, the first evidence for electrical potential differences from bovine strain *E. granulosus* that showed significant differences with previously reported *E. granulosus* from the ovine strain [8]. These results may help understand the molecular mechanisms associated with ion transport and hydroelectrolytic balance into the parasite.

## Materials & Methods

### Parasites

Protoscoleces of *Echinococcus granulosus* were obtained from hydatid cysts of naturally infected cattle lungs from a local abattoir. The parasites were collected and washed as originally described [12] and resuspended in PBS (pH 7.4) supplemented with penicillin (100 units/ml), streptomycin (100 μg/ml), and amphotericin (0.25 μg/ml). The PSCs were kept at 4°C and used up to one month after harvesting. Viability was evaluated based on both the methylene blue exclusion method and microscope examination (10X) of body movements.

### Saline solutions

PSCs were incubated in Ringer Krebs Solution (RKS) containing 121 mM NaCl, 5 mM KCl, 22.5 mM MgCl_2_, 2.5 mM CaCl_2_, 10 mM Hepes and 5.6 mM glucose, pH 7.4. All other constituents, osmolarity, and pH were preserved.

### *In vitro* incubation procedures

Parasites were stored at 4°C in PBS (supplemented with antibiotics and antimycotics) and immediately before the impalement, PSCs were washed and preincubated for 2 h at 37°C in fresh RKS without antibiotics.

### Electrophysiology

Electrical recordings were conducted with a single microelectrode high input impedance (>10^11^ Ω) amplifier-intracellular electrometer (Model IE-210, Warner Instruments, Hamden, CT) with an internal 4-Pole low-pass Bessel filter set at 20 kHz and sampling rate at 10 kHz. The electrometer was connected in parallel to an Analog-Digital Converter (TL-1 interface. Tecmar, Solon, OH) that fed the digital input of a personal computer running Axograph (Axon Instruments, Union City, CA, USA) as a digital oscilloscope.

### Microelectrode recordings

Microelectrodes from glass capillaries (Biocap, Buenos Aires, Argentina) with 1.25 mm internal diameter were pulled on a PB-7 pipette puller and heat-polished on an MF-9 pipette polisher (Narishige, Tokyo, Japan), and filled with filtered 3 M KCl solution. Tip resistance ranged between 10 and 40 MΩ. The reference electrode was a wider tip glass capillary, also filled with 3 M KCl solution, connected to Cl^-^-plated silver wire (Ag/AgCl) and to the ground socket of the electrometer. The recording chamber consisted of a glass slide where an aliquot (500 μl) of a protoscolex suspension was added (~100 PSCs/ml). Parasites were impaled after being individually captured and held with a suction micropipette (Fig. 1c). Impalements were performed under optical microscopy with an IMT2 Olympus inverted microscope (x10). All impalements were performed at room temperature after PSC incubation at 37°C. The criteria for acceptable impalement included: (1) a sharp deflection to a peak potential after electrode penetration into the PSC; and (2) an abrupt return to the original baseline (0 mV) after removal from the parasite.

**Fig. 1:**
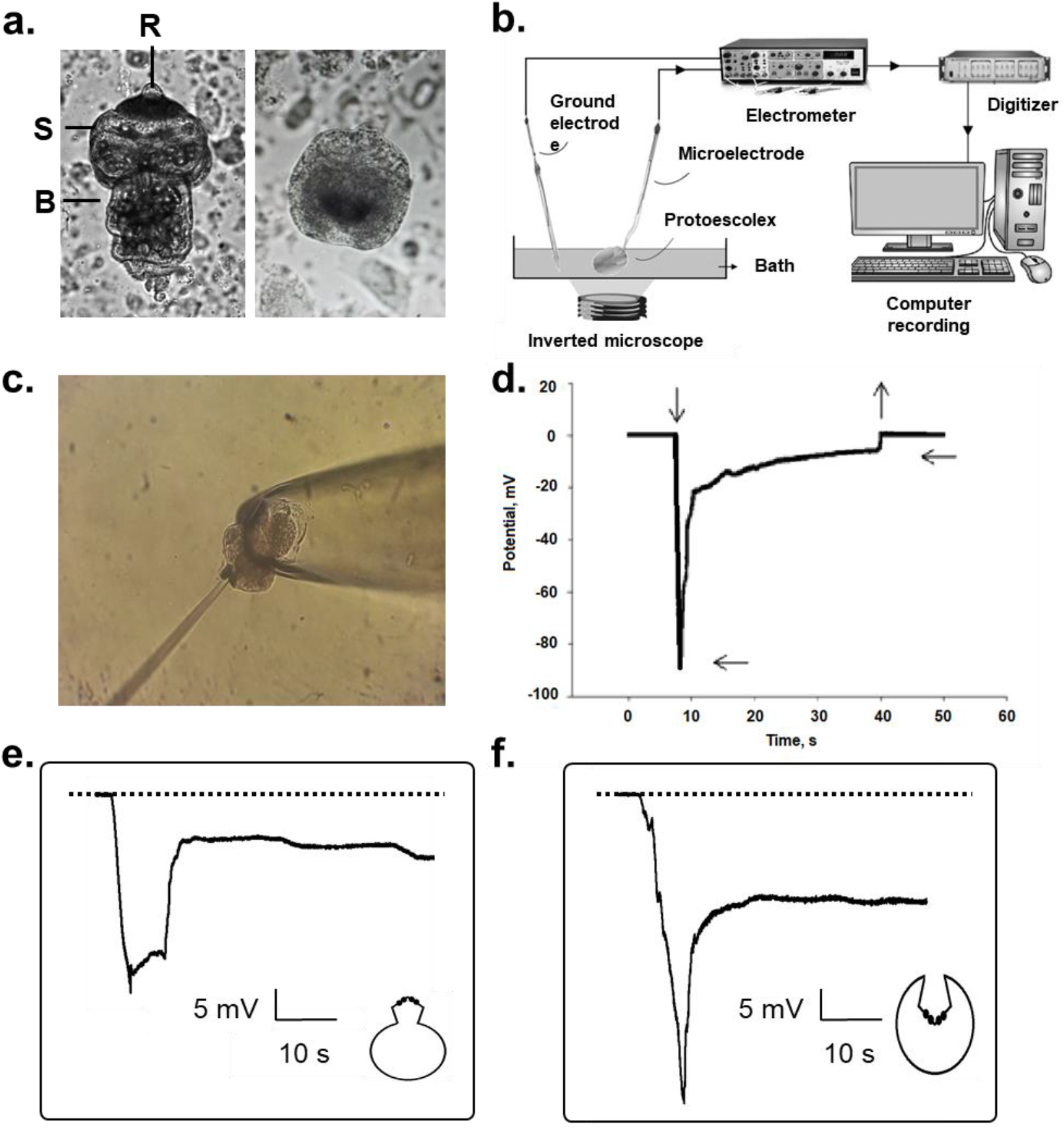
Electrical microelectrode recordings of PSCs from *E. granulosus*. **(a)** Left: Evaginated PSC; **R**= Rostellum, **S**= Sucker, **B**= Body; Right: Invaginated PSC. **(b)** Electrical setup. Both ground and impaling microelectrodes were connected to an electrometer and recorded through an A/D system to a personal computer. **(c)** Evaginated PSC held by a suction pipette and impaled with a recording microelectrode (shown on Left). **(d)** Representative recording of an invaginated PSC shows deflections upon impalement (downward vertical arrow) and withdrawal (upward vertical arrow). A typical tracing shows transient (peak, PD_1_) and more stable lower (plateau, PD_2_) potentials. Horizontal arrows indicate recorded values for PD_1_ and PD_2_, respectively. **(e)** Example of the tegumental potential of invaginated PSC. **(f)** Example of the tegumental potential of evaginated PSC. Dashed lines represent 0 mV. Please note that no upward deflections after withdrawal of microelectrode are indicated.

### Data analyses

Data analysis was conducted with Clampfit 10.7 (Axon Instruments). Sigmaplot 11.0 (Jandel Scientific, Corte Madera, CA) was used for statistical analysis and graphics. Unless otherwise indicated, only values from animals up to 6 days post-harvest were considered for statistical analyses due to significant changes in tegumental voltages afterward. Normal distributions of data were examined using the Shapiro-Wilk W test for normality where *p* < 0.05 indicated a significant departure from normality. Upon failure of the Shapiro-Wilk W test, the Box-Cox normality plot was performed to transform data for normalization [13]. Correlation between potential differences (PD) was performed by the linear regression model, testing the resultant equation against the null hypothesis of a slope equal to zero, considering a *p* < 0.05. Student t-test and one-way ANOVA were used to determine statistical significance between different experimental groups. Averages of corrected data values were expressed as the mean ± SEM for each experimental condition per number of (*n*) or per number of PSCs (*N*).

## Results

### Microelectrode recordings of protoscoleces

The electrical behavior of bovine lung PSCs was explored by microelectrode impalement as previously reported for *Schistosoma mansoni* [9, 14-16] and PSCs of *E. granulosus* of ovine origin [8]. Both invaginated and spontaneously evaginated PSCs were impaled. Upon microelectrode penetration of the **tegumental surface (Fig. 1c) a rapid negative transient deflection in electrical potential was always recorded (*n* = 135, Fig. 1D), which we referred to as “peak” or potential difference 1 (*PD_1_*)**.

This transient potential always spontaneously decayed to another lower, referred to as “plateau” potential (*PD_2_*), even when the microelectrode was not advanced more deeply into the parasite. Both *PD_1_* and *PD_2_* were always discernible and observed in either invaginated or evaginated parasites. Although impalements for either invaginated or evaginated PSCs were always conducted under similar conditions, raw values did not follow a Normal distribution (Fig. 2). Thus, group values (either invaginated or evaginated PSCs) were transformed by the Box-Cox algorithm [13], to allow quantitative comparisons. Relative to the bath (0 mV), the *PD*_1_ values for invaginated PSCs ranged between −179.4 and −14.4 mV with a mean of −64.8 ± 1.4 mV (*n* = 49). The *PD*_2_ values for the same group ranged from −88.4 to −6 mV, with a mean value of - 23.2 ± 0.3 mV (*n* = 49), thus representing a Δ*PD* (*PD*_1_ – *PD*_2_) of −41.6 mV (Table 1). Evaginated parasites instead, had *PD_1_* values that ranged from −231.3 to −20.7 mV with a mean of −92.9 ± 0.5 mV (*n* = 86). These potentials shifted to a *PD*_2_ value that ranged between −77.5 mV and −23.6 mV, with a mean value of −33.5 ± 1.8 mV (*n* = 86), representing a Δ*PD* (*PD*_1_ – *PD*_2_) of −59.4 mV (Table 1). Thus, mean *PD*_1_ and *PD*_2_ values were statistically different among groups, indicating different tegumental electrical properties (*p* ≤ 0.05; Table 1). The Δ*PD* values were also statistically different among groups (Table 1), being more negative in the evaginated state (*p* < 0.0001).

**Fig. 2:**
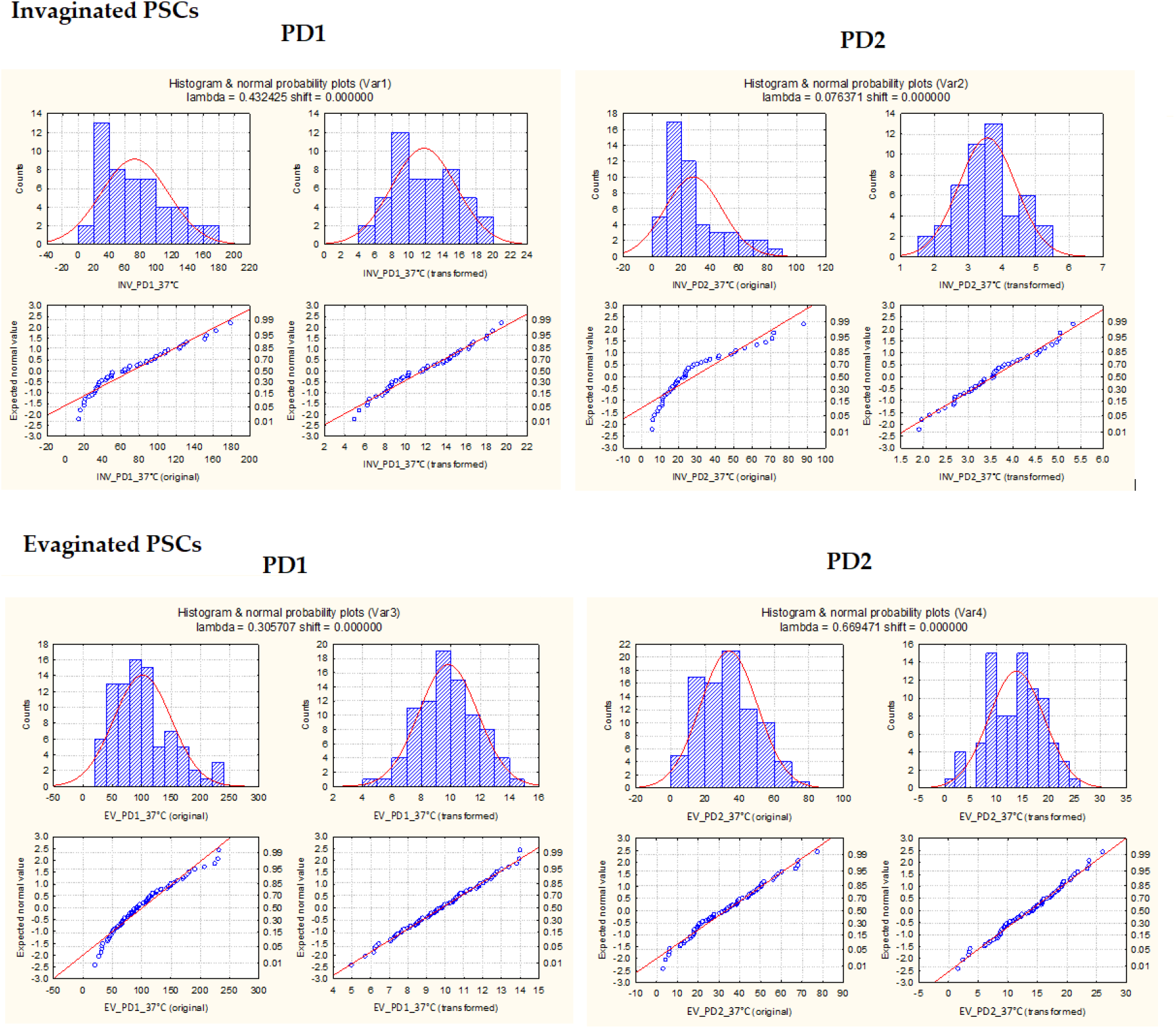
Tegumental potential data distributions. Recordings of PSCs from *E. granulosus*. Left panels indicate PD_1_ values and right panels PD2 values, for invaginated (Top panels, n = 49 values) and evaginated PSCs (Bottom panels, n = 86 values), respectively. Values (mV) are expressed as the mean ± SEM for peak (*PD*_1_) and plateau (*PD*_2_), respectively.

**Table 1.**
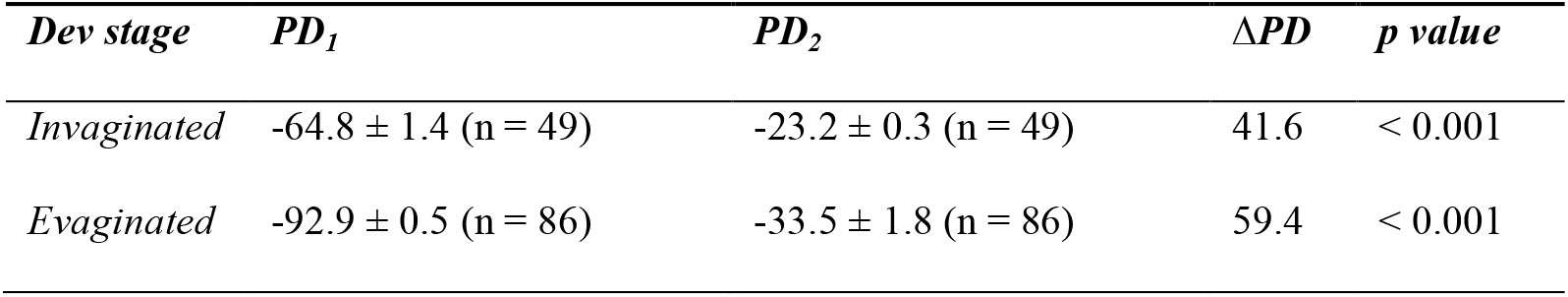
Tegumental potentials of PSCs from *E. granulosus*.

To explore whether impaling itself modified the PD values by either tissue damage or KCl leakage from the pipette, multiple sequential measurements (five events/organism) were also conducted in individual PSCs. For invaginated PSCs, mean *PD_1_* values were - 79.3 mV ± 11.5 (*N* = 15) that decayed to a mean *PD_2_* value of −32.6 mV ± 4.9 (*N* = 15). For evaginated PSCs, *PD_1_* was −93.1 mV ± 14.3 (*N* = 30) that decayed to a *PD_2_* value of −35.3 mV ± 7.2 (*N* = 30). Thus, mean individual values did not differ significantly from those obtained from a large number of individual measurements (*p* > 0.05; Table 1), and therefore impalement itself did not contribute to the transient peaks (*PD*_1_), nor affected the Δ*PD* values observed in either developmental stage of the parasite.

### Time response of the impalements

Because of the different enfolding in the tegumental epithelium between the invaginated and evaginated parasites, the time taken for full peak polarization after impalement was also compared between groups. Depolarizations lasted between 100 and 1000 milliseconds, with slopes of −3.7 ± 0.02 mV/sec (*n* = 8), and −10.3 ± 0.13 mV/sec (*n* = 5) for invaginated and evaginated PSCs, respectively. Thus, the change in potential at impalement was three times faster in evaginated PSCs (*p* < 0.001). The *PD_1_* and *PD*_2_ potentials were well defined and statistically different within and among groups, as determined in both invaginated and evaginated PSCs (*p* < 0.001, Table 1). However, microelectrode recordings were statistically higher in evaginated PSCs than invaginated PSCs, representing a Δ*PD*_1_ of almost 30 mV, and Δ*PD*_2_ of only 10 mV (*p* < 0.05, Table 1). To explore whether the developmental stage of evagination affected this interaction, the correlation between *PD*_1_ and *PD*_2_ was also determined in either stage (Fig. 3c and 3d). A statistically significant correlation was observed between the magnitude of the tegumental and intra-parasitic potentials (*r* > 0.5), with a slope of 0.199 ± 0.029 (*n* = 49) and 0.266 ± 0.054 (*n* = 86) for invaginated and evaginated PSCs, respectively. However, the correlations were not significantly different among groups, suggesting electrical continuity between the functional compartments in both invaginated and evaginated PSCs.

**Fig. 3:**
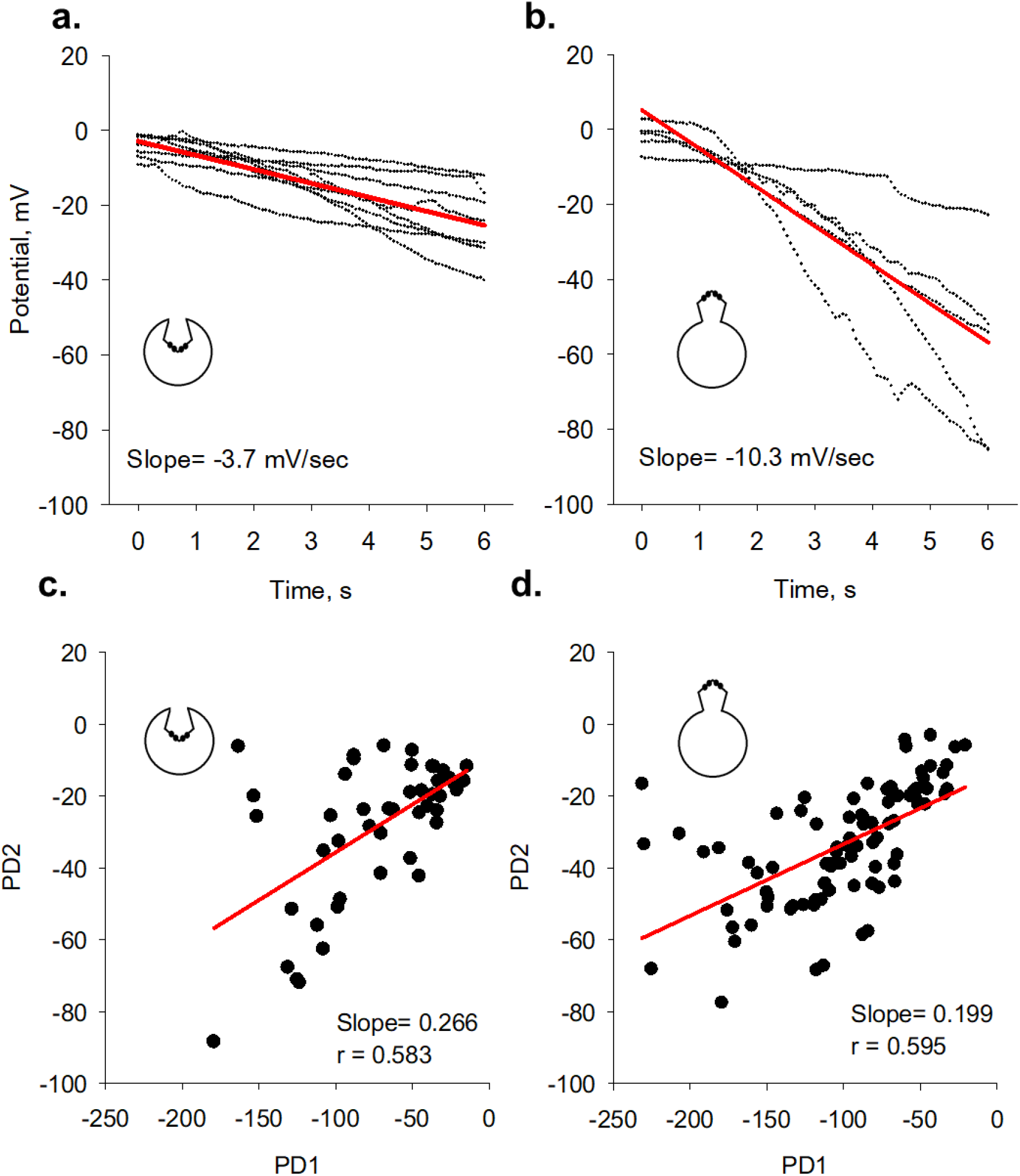
Time-course of impalement and correlations between PD_1_ and PD_2_. Impalements were carried out at 37°C in RKS, for both invaginated and evaginated PSCs (Left and Right panels, respectively). Solid red lines indicate the best linear fitting of time deflections (shown as dotted lines) under each condition. Plots **(a)** & **(b)** show slopes in mV/sec as shown at bottom. Plots **(c)** & **(d)** show the correlation between PD_1_ (mV) and PD_2_ (mV), for invaginated and evaginated PSCs, respectively.

### Effect of aging on the tegumental potentials

Electrical recordings varied through time post-harvesting and decreased significantly after the sixth day for invaginated PSCs and after the twelfth day for evaginated PSCs (Fig. 4). Therefore, mean values were considered from measurements conducted up to a week after collection from hydatid cysts.

**Fig. 4.**
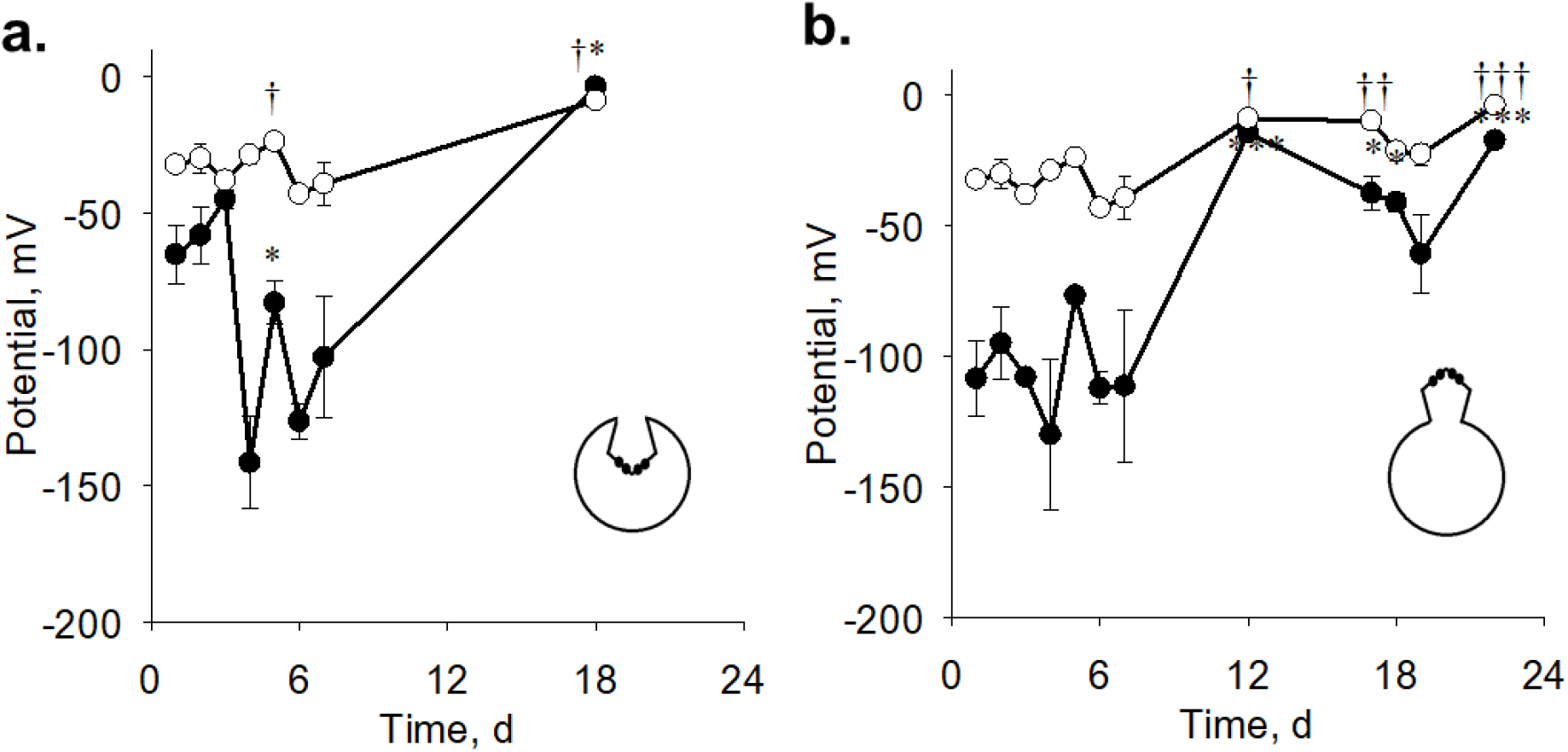
Effect of aging on the electrical potentials of PSCs from *E. granulosus*. Graphs show parasitic potentials at different times post-harvest in days (d) from invaginated **(a)** and evaginated PSCs **(b)**, respectively. Filled symbols correspond to PD_1_ and open symbols to PD_2_. Values are the mean ± SEM, for *n* between 3 and 56. Symbols † or * indicate statistically significant difference from day 1 (PSCs harvesting) for PD_1_ or PD_2_ respectively, with †p < 0.05, ††p < 0.01, †††p < 0.001 and *p < 0.05, ***p < 0.001.

### Parasitic potentials in identified anatomical regions of evaginated PSCs

While invaginated PSCs may be expected to represent a single parasitic compartment, evaginated parasites may offer distinct and well-defined anatomical regions [17, 18]. Because similar correlations were observed between *PD*_1_ and *PD*_2_ in both populations (Fig. 3c and 3d) we also explored whether the process of evagination exposed a particular electrical compartment not defined in the invaginated state. For this, recordings were obtained from the three identifiable anatomical regions of evaginated PSCs [17, 18] including the rostellum, the sucker, and body regions of the parasite (Fig. 1a). *PD*_1_ and *PD*_2_ remained different among regions, with Δ*PD* in the 36.8 to 50.7 mV range (Table 2). The trans-tegumental potential (*PD*_1_) obtained from the rostellum was significantly higher than that from either the neck or body of the evaginated PSC (Table 2). Even, intra-parasitic potential (*PD*_2_) showed significant statistical differences among the three anatomical regions, suggesting the distinct compartmental distribution of potential throughout the parasite.

**Table 2.** Tegumental potentials from different anatomical regions of evaginated PSCs.

| | *n* | *PD1* | *PD2* | $\Delta PD$ | *p value* |
| --- | --- | --- | --- | --- | --- |
| ***Rostellum*** | 20 | $-81.5 \pm 1.4^a$ | $-30.8 \pm 0.8^a$ | 50.7 | < 0.001 |
| ***Sucker*** | 64 | $-65.9 \pm 2.2^b$ | $-27.4 \pm 0.7^b$ | 38.5 | < 0.001 |
| ***Body*** | 42 | $-60.4 \pm 1.8^b$ | $-23.6 \pm 0.7^c$ | 36.8 | < 0.001 |

Mean ± SEM followed by the same letter in each column were not significantly different from each other (Tukey's test, $p < 0.01$ ).

## Discussion

The present study provides, to our knowledge, the first reported characterization of the tegumental electrical properties of PSCs of *Echinococcus granulosus* from bovine lung. The values and electrical features were different from those reported for PSCs of *Echinococcus granulosus* from the ovine strain [8] and interestingly were more similar to those of *Schistosoma mansoni* [9]. We observed that upon penetration of the tegumental surface of the PSC, the first peak of negative potential (*PD*_1_) was recorded that was lower in invaginated PSCs as compared to evaginated PSCs. The *PD*_1_ values were independent of the duration or depth of the impalement, and spontaneously decayed to a lower plateau value (*PD*_2_), which remained stable and, in all cases was lower in invaginated PSCs.

Because the syncytial tegument of *E. granulosus* is very thin, 2 to 3 μm wide [19], and despite the actual location of the impaled microelectrode was not ascertained, the first peak voltage deflection (*PD*_1_) should correspond to the trans-tegumental potential, in agreement with previous reports of ovine PSCs of *E. granulosus* [8], and *Schistosoma mansoni* [9, 20]. Elimination of the tegument by addition of either deoxycholate or Triton X-100 always elicited a rapid and irreversible depolarization of this particular electrical potential difference. Moreover, it was demonstrated by iontophoretic injection of horseradish peroxidase into *S. mansoni* that *PD*_1_ originated in the tegumental epithelium [20]. The second potential observed in that study, which is similar to *PD*_2_ obtained in the present report, was ascribed to muscle masses below the tegumental membrane [9, 20]. This is also in agreement with electrophysiological studies describing the degree of electrical coupling between tegument and muscle of *S. mansoni*, were presented by Thompson et al [16].

Aging is another parameter that may bring information regarding the location of *PD*_1_ and *PD*_2_. Ibarra and Reisin [8] reported that under control conditions, *E. granulosus* PSCs from ovine origin remained stable up to 25 days. However, the present study on PSCs from bovine lung origin showed a significant decrease in *PD*_1_ after 6 days post-harvest, as compared to *PD*_2_. On the other hand, no invaginated PSCs could be found after 20 days post-harvest, because gradual development into the evaginated stage.

The reason(s) for the differences between the shape and magnitude of the PDs in ours and Ibarra and Reisin [8] could span from technical details of the electrical recordings to the origin and species of the PSCs, including the *E. granulosus* genotypes. Further experimentation will be required to ascertain the nature of these changes.

The present study indicates that although the general properties of the syncytial tegument may remain the same in different developmental stages of the parasite, the speed of depolarization at impalement and *PD*_1_ and *PD*_2_ values were statistically higher in the evaginated stage as compared to the invaginated stage of the parasite. Thus, it is possible that the invaginated parasite could provide a second electrical barrier that is eliminated upon evagination. However, the magnitude of *PD*_1_ and *PD*_2_ were similarly correlated in both developmental stages, an indication that the electrical compartments remained identically coupled.

Previous studies have not reported on the tegumental potentials of the evaginated stage of the PSC from *Echinococcus granulosus.* The electrical data of the anatomical regions of the evaginated parasite obtained in the present study further suggest that different parts may provide specific contributions to the invaginated stage, where the anatomical regions such as the rostellum and the neck are intra-parasitic. Moreover, different tegumental potentials have been reported in anatomical regions of *Schistosoma* [16].

Little is known as to the electrodiffusional pathways that contribute to the electrical potential differences in *Echinococcus* and other flatworms. Cantiello, Ibarra & Reisin [21] observed that the passive K^+^ influx corresponded to a simple diffusional mechanism distributed in at least two compartments, one small and faster and the other large and slower ionic exchange, although no anatomical correlates of the parasite were reported. The ion channel species responsible for these movements have yet to be identified, as well as their contribution to the electrophysiology of the parasite. Grosman and Reisin provided preliminary evidence for the presence of cation-selective channels in extracted membranes of *E. granulosus* PSCs from ovine origin [22, 23]. The recent genomic sequencing of *E. granulosus* and other cestodes [6, 24] may help identify the molecular fingerprints of the channel and transporter species, and assess their relevance as possible pharmacological targets [25]. To date, useful antiparasitic drugs include potential Ca^2+^ channel blockers such as praziquantel [26, 27], and benzimidazoles, such as mebendazole and albendazole, that modify the microtubular cytoskeleton [28]. New ion channel targets and relevant interactions could be expected to bring about novel therapeutic strategies [29-31]. We recently obtained preliminary information to suggest that known links between excitable cation channels and the actin cytoskeleton [32] may help potentiate the ability of praziquantel to paralyze the *Echinococcus granulosus* PSCs [33].

In summary, the present study provides evidence that PSCs of *Echinococcus granulosus* from the bovine strain present complex tegumental potentials remarkably differ from those previously reported for a similar preparation from sheep origin. Two distinct functional compartments were identified of tegumental origin and extracellular domain of the intra-parasitic milieu, respectively. Future studies will be required to assess the nature of the differences, genetic or otherwise from those reported with ovine *E. granulosus*, and those expected from other intermediary species. The microelectrode measurements of parasitic potentials may prove an invaluable tool in helping characterize the contribution of various ion channel species and enzymatic transporters in cestodes, and also provide a rapid testing approach to explore and identify new pharmacological targets.

## Acknowledgments

The authors wish to acknowledge the National Research Council of Argentina (CONICET) for funding of the present study (PUE 2292180100054CO).

## Author Contributions

**Conceptualization:** Horacio F. Cantiello

**Data curation:** Mónica P. A. Carabajal, Marcos A. Durán, Santiago Olivera

**Resources:** Santiago Olivera, María José Fernández Salom

**Formal analysis:** Mónica P. A. Carabajal, Horacio F. Cantiello

**Funding acquisition:** Horacio F. Cantiello

**Supervision:** Horacio F. Cantiello

**Investigation:** Horacio F. Cantiello, Mónica P. A. Carabajal, Marcos A. Durán, Santiago Olivera, María José Fernández Salom

**Writing – original draft:** Horacio F. Cantiello, Mónica P. A. Carabajal

**Writing – review & editing:** Horacio F. Cantiello

